# Nuclear export of the pre-60S ribosomal subunit through single nuclear pores observed in real time

**DOI:** 10.1101/2021.03.30.437662

**Authors:** Jan Andreas Ruland, Annika Marie Krüger, Kerstin Dörner, Rohan Bhatia, Sabine Wirths, Daniel Pòetes, Ulrike Kutay, Jan Peter Siebrasse, Ulrich Kubitscheck

## Abstract

Ribosomal subunit biogenesis within mammalian cells initiates in the nucleolus with the assembly of a 90S precursor particle, which is subsequently split into the pre-40S and pre-60S subunits. During further processing steps, pre-ribosomal subunits are loaded with export receptors, which enables their passage through the pore complexes (NPCs) into the cytoplasm. Here export factors are released and both subunits can form a mature ribosome. Ribosomal biogenesis has been studied in great detail by biochemical, genetic and electron microscopic approaches, however, until now live cell data on the in vivo kinetics are still missing.

We analysed export kinetics of the large ribosomal subunit (“pre-60S particle”) through single NPCs in living human cells. To assess the in vivo dynamics of this process, we established a stable cell line co-expressing Halo-tagged eIF6 and GFP-fused NTF2 to simultaneously label ribosomal 60S subunits (eIF6) and NPCs (NTF2). By combining single molecule tracking and super resolution confocal microscopy in a highly customized microscopic setup, we visualized the dynamics of single pre-60S particles during the interaction with and export through single NPCs. In this way we obtained unprecedented insights into this key cellular process.

Our results revealed that for export events, maximum particle accumulation is found in the centre of the pore, while unsuccessful export terminates within the nuclear basket. The export process takes place with a single rate limiting step and an export dwell time of **~**24 milliseconds. Only about 1/3 of attempted export events were successful. Given the molecular mass of the pre-60S particles our results show that the mass flux through a single NPC can reach up to ~125 MDa·s^−1^ in vivo.

## Introduction

Ribosome biosynthesis is a cellular mammoth task requiring the orchestrated interplay of more than 200 proteins over three different compartments in the eukaryotic cell. Mature ribosomes contain four ribosomal RNAs (rRNA) and ~80 proteins and are built from the small 40S and the large 60S subunit. The biogenesis of the subunits is a complex process and begins with the synthesis of the rRNAs, which provide the backbone for both subunits (for review see ^1,2^). In eukaryotic cells, three of the four rRNAs are transcribed by RNA polymerase I (Pol I) as one single precursor in the nucleolus, while the 5S rRNA is transcribed by Pol III. Pre-rRNA synthesis, cleavage and maturation go along with the integration of ribosomal proteins and of multiple accessory factors and finally results in the small pre-40S and a large pre-60S subunit. The eukaryotic pre-60S particle (MW 2.1-3.1 MDa)^3,4^ is about 25 nm in diameter while the pre-40S subunit has an average diameter of 16 nm. The subunits are among the bulkiest RNA transport substrates that have to be exported out of the nucleus.

The export pathways of the different RNA classes can be distinguished by the transport receptors used and how they are loaded to the RNA (see^5–7^ and refs. therein). Small RNA molecules like transfer RNA, micro RNA or small nuclear RNAs are exported by ‘importin β-like’ receptors, e.g. Exportin t or Exportin 5, which are bound directly in a RanGTP-dependent manner. In contrast, rRNA and messenger RNA (mRNA) are exported as intricate ribonucleoprotein (RNP) particles. The export of mRNA requires Mex67/Mtr2 in yeast and Tap/p15 (or NXF1/Nxt1) in human and their mRNP incorporation is mediated by the adaptor proteins. The directionality of the mRNA export does not rely on the RanGTP/GDP gradient. Rather, an ATP-dependent RNA helicase, DDX19 in mammals (Dpb5 in yeast), together with additional factors remodels and releases the mRNPs after export into the cytoplasm^8,9^.

In yeast Mex67/Mtr2 is also involved in the export of the ribosomal subunits^10^ but acts in doing so together with additional export factors, i.e. Rrp12p^11^, Arx1^12,13^, Ecm1^14^, Bud20^15^, Npl3^16^ and Gle2^17^ for the pre-60S particle (for review, see^7^). The Mex67/Mtr2 heterodimer is bound to the late pre-60S subunit by direct interaction of its loop insertions in the middle NTF2-like domains of both Mex67 and Mtr2 with the 5S-rRNA^10^. Tap/p15 does not have these insertions and thus does not play a role in ribosomal export. Instead, in human cells Exportin-5 (Exp5) is also needed for pre-60S export^18^.

Although ribosomes are evolutionary highly conserved the only transport receptor unambiguously identified in the nuclear export of both the yeast and the human pre-60S particle is Xpo1/Crm1, which recognizes the nuclear export signal (NES) of ribosome-bound NMD3 in a RanGTP-dependent manner^19–21^. After export the Crm1 and Exp5 are removed by RanGAP induced GTP hydrolysis on Ran (reviewed in^22^).

The most detailed view of the actual pore transit of pre-60S particles derives from a recent electron tomographic study^23^. These researchers analyzed NPC passage of the pre-60S particle in snap-frozen yeast cells. They did not label the subunits but identified them in transit through NPCs by their characteristic size and morphology. It was shown that the pre-60S particles move through the central channel of the NPC and that roughly 4-5% of all yeast NPCs contain a subunit in transit at any given time point. Of course, EM studies can only provide a static view of a dynamic process but using a skillful probabilistic approach Delavoie et al. estimated a translocation time of approximately 90 ms.

So far live cell data on the ribosomal export are missing. However, pre-40S and pre-60S particles represent large RNP particles that can be labeled by a comparable approach as we have used previously for transport experiments on mRNA particles^24^. Only recently we succeeded in labeling the pre-40S particles using DIM2 (Pno1) coupled C-terminally to a SnapTag^25^. Our approach enabled us to observe pre-40S particles inside cell nuclei at the single particle level. Here, we used a similar approach to visualize pre-60S particles by employing eIF6, which is loaded on to the pre-60S ribosomal subunit in the nucleolus. Already some years ago it was demonstrated that eIF6-HaloTag is functionally incorporated into pre-60S particles and represents a feasible approach to fluorescently label pre-60S particles in vivo^26^. Here, we established a HeLa cell line stably expressing eGFP-tagged nuclear transport factor 2 (NTF2)^27^, a transport receptor highly enriched within NPCs in order to precisely localize single nuclear pore complexes. This cell line was re-transduced to additionally express eIF6-HaloTag, which was subsequently labeled with the membrane permeable dye JF549^28^. By adding JF549 at sub-nanomolar concentrations we succeeded to optically singularize pre-60S particles, whose trajectories could well be traced by single particle tracking. The quasi-simultaneous observation of single NPCs by super resolution confocal laser scanning microscopy enabled the real-time observation of nuclear pre-60S particle export. From the single particle data we could deduce the export dwell time, locate the route of successful and unsuccessful export events and identify a rate limiting step.

## Results

### eIF6-Halo tagged pre-60S particles are functional

In order to label the pre-60S particles, we targeted eIF6, which is stably incorporated into the particle^29^ and prevents its premature association with the pre-40S subunit. We created a HeLa cell line stably expressing both eGFP-NTF2 and eIF6-HaloTag. The HaloTag allows for labelling with photostable dyes such as JF549 and to visualize single pre-60S particles by adjusting the dye concentration. Staining eIF6-HaloTag by JF549 at a concentration of 0.1 μM resulted in the typical nucleolar accumulation of ribosomal subunits (Fig 1A and supplemental Figure 1) and suggested that the Halo-tagged eIF6 exists preferentially bound to pre-60S particles as was previously concluded by Gallo and coworkers^26^.

**Fig. 1.**
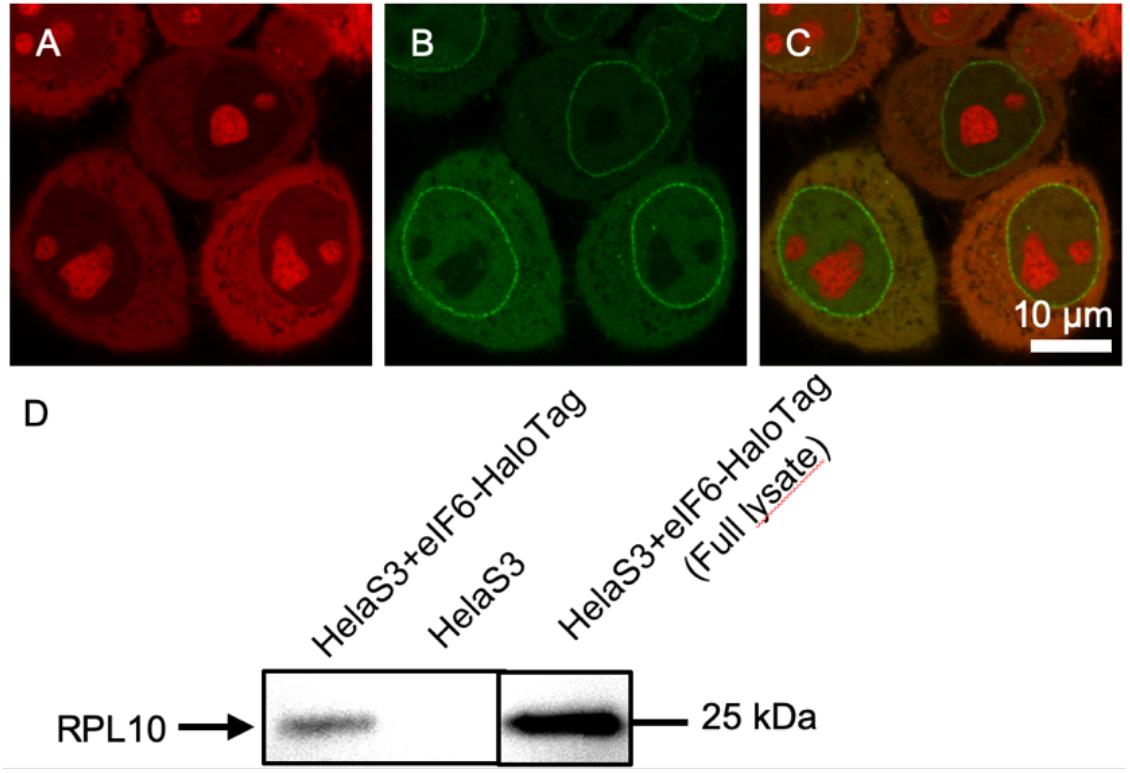
HeLa S3 cells expressing eGFP-NTF2 and eIF6-HaloTag. (A) eIF6-HaloTag-JF549 staining. (B) eGFP-NTF2. (C) Overlay of both channels. (D) Immunoblotting against Rpl10 after pulldown of eIF6-HaloTag in HeLa S3 cells stably expressing eIF6-HaloTag and untreated HeLa S3 cells. Full lysate is shown as antibody control.

To further test whether the tagged eIF6 acts like the wild type protein, we performed pulldown experiments using the HaloTag. Immunoblotting revealed that the purified complex contained Rpl10 and confirmed that the tagged eIF6 is successfully assembled into the subunit.

The biochemical and microscopic data were also verified when we treated the eIF6-JF549 cells with the transcriptional inhibitor Actinomycin D, which resulted in a complete loss of nucleolar labelling (supplemental Figure 2).

To test if labelled pre-60S particles can be exported successfully, we treated the cells with Leptomycin B (LMB). LMB prevents binding of the export receptor Crm1 to NMD3 and thus blocks the nuclear export of pre-60S particles^20,30^. Correspondingly, we observed an increase in intranuclear fluorescence when we applied LMB to HeLa cells expressing eIF6-HaloTag molecules, which were marked by JF549 (supplemental Figure 3). The fluorescence increase was clearly detectable, but not exhaustive. We suspect this was due to a fraction of unbound eIF6-HaloTag-JF549 molecules that were still able to shuttle across the NE in this bulk experiment.

### Tracking of single pre-60S particles at single NPCs

By adjusting the JF549 concentration between 0.3 and 1 nM we could visualize single pre-60S particles in live cell measurements. In order to directly observe the intracellular transport of single pre-60S particles we combined a narrow-field HILO illumination^31^ and EMCCD detection with an LSM 880 Airyscan in a customized microscopic set up. This enabled us to acquire movies of single pre-60S particles using the EMCCD detection path and parallel detection of GFP-tagged NPCs revealing the position of the nuclear envelope (NE) in the very same cell using confocal Airyscan imaging (see Fig. 2 and movie S1). The particles were mostly immobile inside nucleoli, but in the nucleoplasm diffusive motion was prevailing. Tracking of the single particles revealed a maximal nuclear diffusion coefficient of 1.7± 0.1 μm^2^/s (see Methods). This value was smaller than the maximal diffusion coefficient that we recently determined for single pre-40S particles, 2.3 ± 0.3 μm^2^/s ^25^, and significantly smaller than the maximal diffusion coefficient of free, unbound proteins in cell nuclei, >10 μm^2^/s ^32^. Together with the results of the bulk experiments reported above, this showed that the respective particles corresponded to single pre-60S particles.

**Fig. 2.**
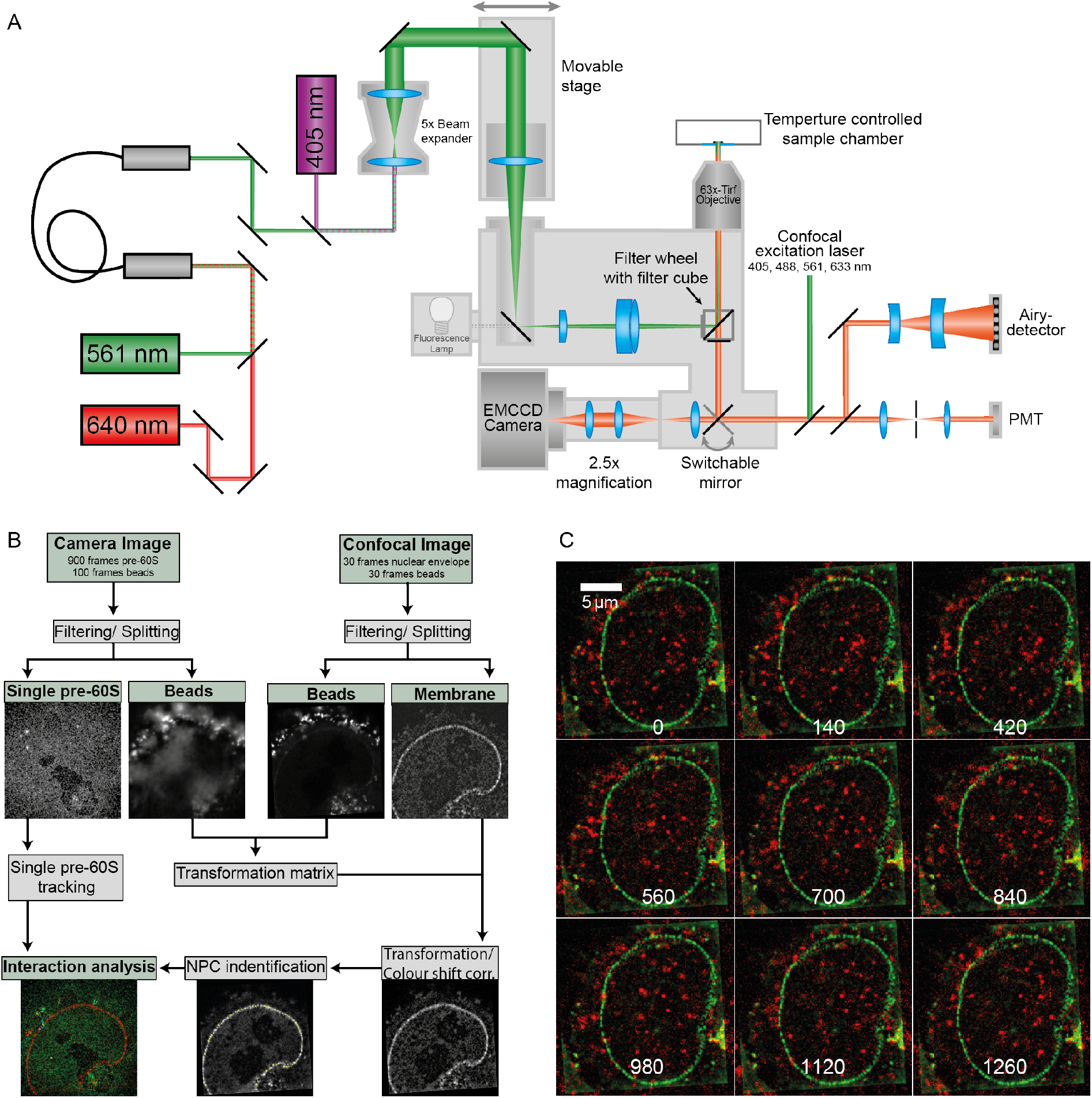
Single molecule and super resolution confocal microscopy of the same cell. (A) A Zeiss LSM880 was extended by an additional beam path for narrow-field single molecule excitation and high sensitivity detection. This beam path was used for imaging single pre-60S particles labelled by eIF6-HaloTag-JF549. By switching the fully automated microscope to confocal super resolution mode we were able to image the very same sample region by Airyscan microscopy, which revealed single NPCs lined up at the NE labelled by eGFP-NTF2. For image registration, UV fluorescent microbeads coupled to the cell surface, which were seen in both modes, were used. (B) Sketch of the measurement principle. Using laser illumination with a wavelength of 561 nm a movie of single eIF6-HaloTag-JF549 inside a cell nucleus was acquired at high frame rate. After movie acquisition illumination switched from 561 to 405 nm to acquire images of the reference beads. Subsequently, super resolution imaging of the NPCs labelled by eGFP-NTF2 and of the reference beads was achieved by Airyscan microscopy in the green and UV fluorescence channels, respectively. The EMCCD movie was used to determine pre-60S particle tracks in the nucleus and across the NE. The super resolution images of the NTF2-labelled NPCs were employed to identify single NPCs and their positions along the NE. The two sets of the reference bead images were used to calculate the transformation matrix that was needed to map the pre-60S trajectories onto the NPC positions. (C) Hela cells stably expressing NTF2-eGFP and eIF6-HaloTag labelled by JF549 after image registration. Single pre-60S particles could be discerned. Numbers represent time in ms.

HeLa cells comprise > 3000 NPCs in their NE^33–35^, and display nearest neighbour distances of about 130 nm ^36^. Our LSM 880 Airyscan achieved optical super resolution (about 150 nm laterally) in the green fluorescence channel using an 63x NA 1.46 objective lens. Thus, with this instrument we were able in many cases to identify single NPCs (Figs. 2 and 3).

**Fig. 3.**
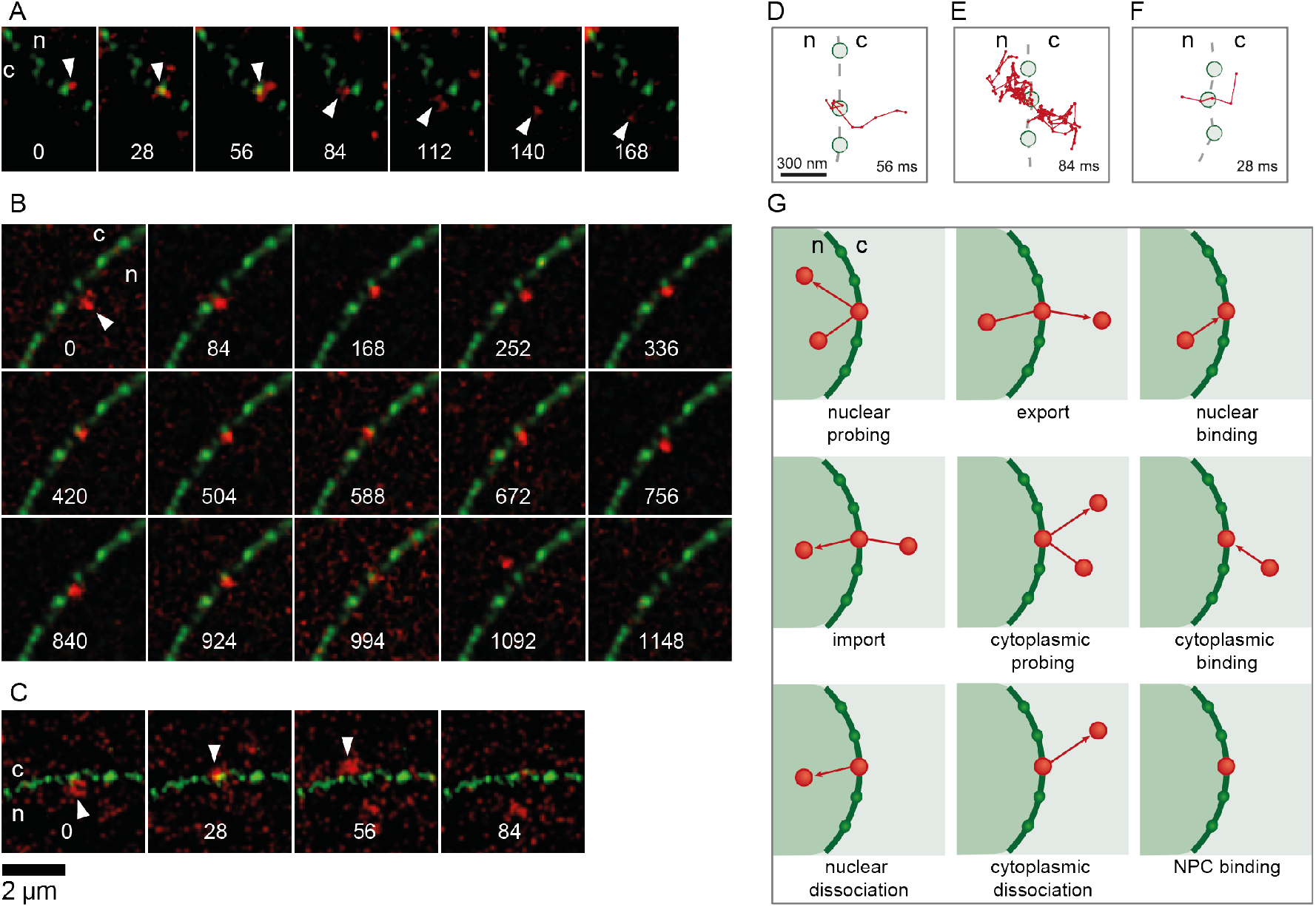
Single pre-60S particles passing a single NPC. (A, B, C) Image sequences showing export events. The numbers represent time in ms. (D, E, F) Trajectories of the particles shown in (A-C) rotated such that the transport direction is parallel to the x-axis, n, nucleoplasm; c, cytoplasm. Also, the position of the central and two neighbouring NPCs were shown. The given times correspond to the respective export dwell times. (G) All possible ways, in which the pre-60S particles could interact with the NPCs.

In order to correlate the observed pre-60S particle positions to locations within the cell nuclei and especially the NE, a precise alignment of the images acquired with the two different detectors was compulsory. To this end we used UV fluorescent microbeads, which were immobilized on the cell surface of the HeLa cells (see Methods). They served as reference markers that could be observed by both the EMCCD and the LSM880. The corresponding point signals were used to register the two fluorescence channels to each other (Fig. 2B and supplemental Fig. 4).

After the image registration process, we could observe single pre-60S particles diffusing within cell nuclei, the NE of which was specified by a line of eGFP-NTF2 loaded NPCs (Fig. 2C). Single subunits could be seen approaching and touching single NPCs. Often, we observed subunits probing one or several NPCs – until finally they were exported into the cytoplasm (Fig. 3A-F, movie S2). In many cases, however, the contact with an NPC did not result in an export, but particles rather returned into the nucleoplasm.

We defined a subunit as interacting with an NPC, when the distance between the subunit location and the NPC position was smaller or equal than a certain threshold, d_int_. The value for d_int_ was deduced from the geometrical extension of the NPC and the localization precision of the respective detection paths, and their combined co-localization precision. This resulted in d_int_ = 129 nm (for details, see Methods).

In order to achieve a quantitative evaluation of the pre-60S particle-NE interactions from the tracking data and the transformed membrane image of each cell we developed a fully automated analysis pipeline comprising three major sections. First, the position of single NPCs in the green channel was automatically determined (see supplemental Fig. 5) returning a list with coordinates of single NPCs. Next, NPC coordinates were compared with the single particle tracking results. Thus, we identified for every individual NPC all tracks of interacting pre-60S particles (see supplemental Fig. 6). We would like to stress that the trajectories identified in this manner did not correspond to all particles, which interacted with the NE, because the NPC list contained only those NPCs, which were clearly singularized and thus identifiable, i.e. were located sufficiently far away from other NPC signals.

Next, the list containing the NPC coordinates was used to create a polygon, which roughly described the shape of the cell nucleus (see supplemental Fig. 7). This polygon was used by a final algorithm that defined the exact type of interaction of the particle with the NPC, i.e. to decide about, e.g., rejected or successful transport events and the direction of transport (Fig. 3G, supplemental Fig. 8). Often, not all steps of the interaction with the NPC could be recorded, e.g. because the observed particle came from or vanished into out-of-focus regions. All observations of interactions with NPCs were sorted into categories by discriminating among all possible interaction types for particles moving near the NE (Fig. 3G). In this manner movies from 230 different cells with an overall run time of almost 90 minutes were evaluated.

### Nuclear export of pre-60S particles

Altogether, we identified 78 complete export processes, for which we could see the particle approaching from the nucleoplasm, interacting with an NPC, and finally dissociating into the cytoplasm. For each export trajectory the interaction duration with the translocating NPC was calculated. A histogram of the translocation times is shown in Fig. 4A. From all export events, we determined a mean translocation time of 35 ± 20 ms. The export dwell time kinetics were obtained by fitting an exponential decay model to the probability distribution of the individual translocation times. To test if different populations of particles with different translocation times exist, the dwell time distributions were modelled by single, double or triple-modal functions (for details, see Methods). The Akaike Information Criterion^37,38^ was used to evaluate the three models. This yielded unambiguously that only a single particle species with a single translocation duration existed. The analysis indicated the existence of a single rate-limiting step. In this manner we determined **τ = 24±4** ms as expectation value for the export duration.

**Fig. 4.**
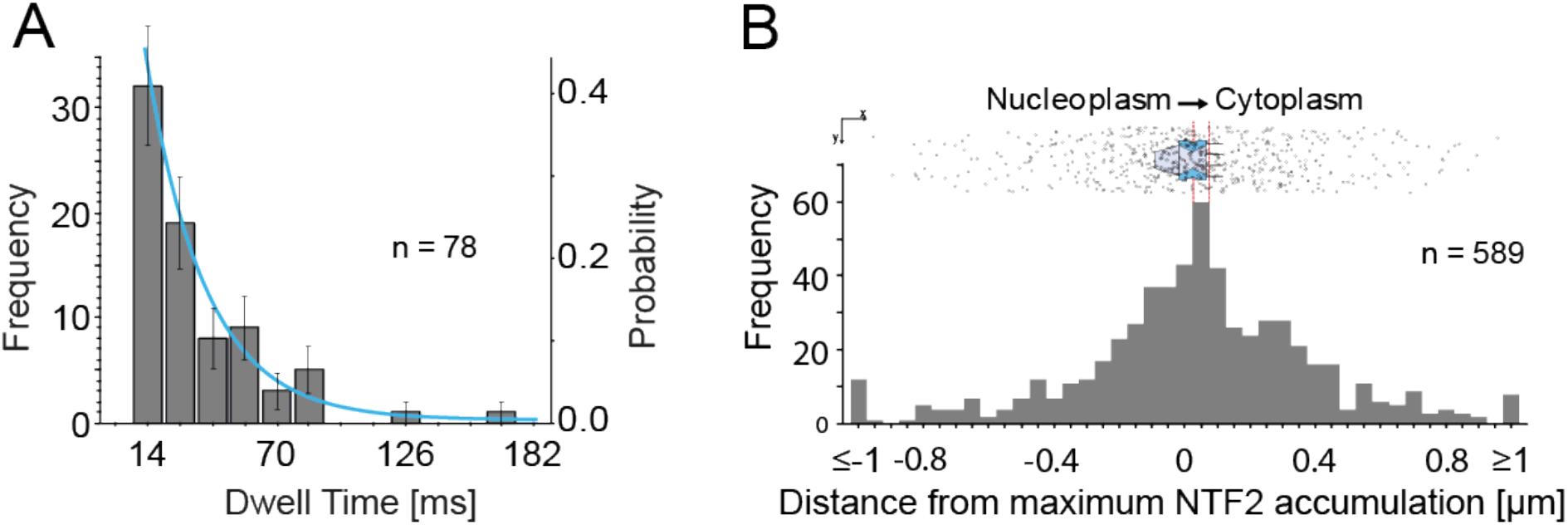
Analysis of pre-60S export events. (A) Distribution of export durations. The shown error bars correspond to the deviation assuming a Poisson process. The blue line indicates the best fit of the probability distribution using an exponential decay model with one translocating species yielding an export duration of τ=24±4 ms. (B) Superposition of all export trajectories aligned to the transporting NPC. (Top) The trajectories were aligned to the transporting NPC (as indicated by the maximum of the eGFP-NTF2 signal) as origin and then rotated such that the export direction coincided with the positive x-axis. (Bottom) The number of pre-60S particle positions in a y-axis region of ± 100 nm was plotted in dependence on the location with regard to the average NTF2 accumulation. The maximum occurred at the position of the cytoplasmic ring of the NPC (red lines).

In order to extract information on the spatial position of the rate-limiting step of the transport process, we aligned all complete export trajectories such that the position of the transporting NPC was defined as the origin of a coordinate system and the transport direction from nucleoplasm to cytoplasm coincided with the positive x-axis (supplemental Figs. 9 and 10). There are various options to accomplish this, we decided to use the two nearest neighbour NPCs as defining the direction of export (for details, see Methods and supplemental Fig. 10). Then, we plotted all export trajectories into this new coordinate system (Fig. 4B, top).

In order to determine the spatial position of the rate-limiting step of the transport process we divided the transport axis into bins of 50 nm and counted the trajectory positions in the respective bins considering all particles that were observed within ± 100 nm of the pore axis in y-direction. For this analysis, we used the point of maximum NTF2 fluorescence intensity as marker for the NPC location. The location of the maximum NTF2 fluorescence was related to the NPC topology using mAb414 labelling the nucleoporin Nup 62 as reference (supplemental Fig. 9 and Methods). Thereby, we found that NTF2 binds preferentially between 0 and 70 nm off the central NPC plane with its maximum at 24 nm towards the nuclear basket. This binding site distribution was in accordance to previous data ^39–41^.

The resulting binding site distribution of the pre-60S particles and a corresponding sketch of the NPC structure indicate that the translocating particles were most often observed in the region of the cytoplasmic ring of the NPC (Fig 5). We assume that the passage of this position corresponded to the rate limiting step of the transport process.

**Fig. 5.**
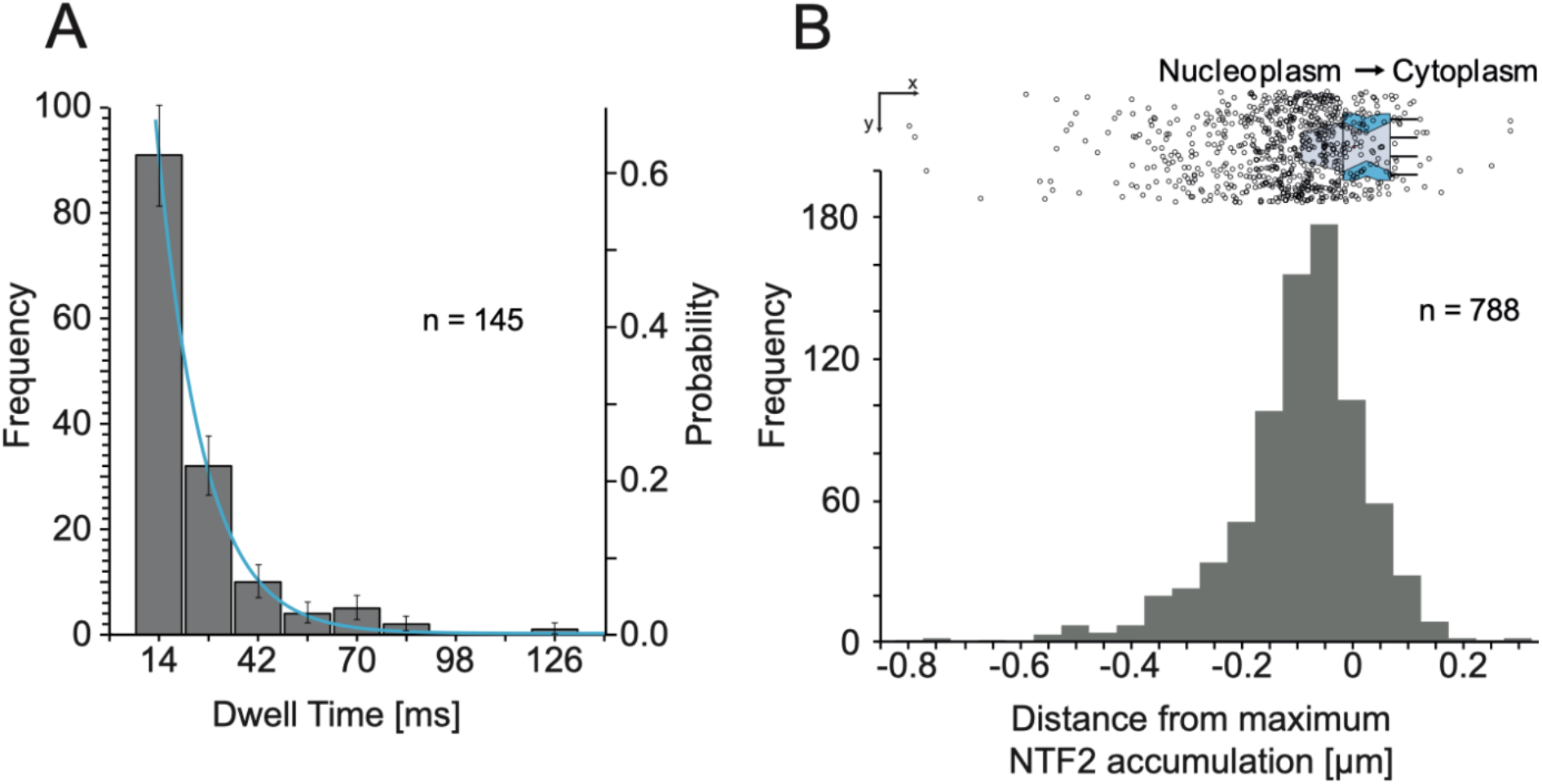
Failed nuclear export of pre-60S particles. (A) Frequency distribution of interaction durations. The shown error bars correspond to the deviation assuming a Poisson process. The blue line indicates the best fit using a single exponential decay function and yielded T=13±3 ms. (B) (Top) The trajectories were aligned to the transporting NPC (as indicated by the maximum of the eGFP-NTF2 signal) as origin and then rotated such that the export direction coincided with the positive x-axis. (Bottom) The number of pre-60S particle positions in a y-axis region ± 100 nm off the central NPC was plotted as a function of the location with regard to the average NTF2 accumulation. The maximum clearly coincides with the nuclear basket.

As mentioned, the analysis pipeline detected nexp=78 reliable export events. This number might seem to be small. However, it was expected that nexp would be relatively low, because a number of strict preconditions had to be fulfilled for an event to be counted. The identification of an export event required that the translocating particle remained in focus during the NE approach, the NPC translocation and dissociation into the cytoplasm. In the 3D space of the cell, diffusive motion away from the focal plane is simply much more probable^24^. Furthermore, our algorithms required a clear and distinct identification of the transporting NPC. In certain cases, we could observe a translocation process, but the corresponding NPC could not unambiguously be identified, e.g. due to a high surface density of NPCs in the respective region of the NE. Additionally, the category “cytoplasmic dissociation” (Fig. 3G and Table 1) contained processes that presumably corresponded to export events, but their first association step with the NPC could not be detected. Similarly, there were numerous approaches to NPCs from the nucleoplasmic side (“nuclear binding”, n=552), for which we did not observe the release from the pore. Presumably, quite a fraction of these resulted in export, whose final step we missed. Also, there were about 800 cases, for which we observed the pre-60S particles only at NPCs and did neither observe the approach nor a release (“NPC binding”). Finally, a valid observation not only required a clearly recognizable line of NPCs in focus and a certain density of particles in the observation field, but also the presence of a sufficient number of UV reference beads in the respective focal observation plane. Fulfilment of all these prerequisites was needed to obtain useful transport data.

**Table 1.**
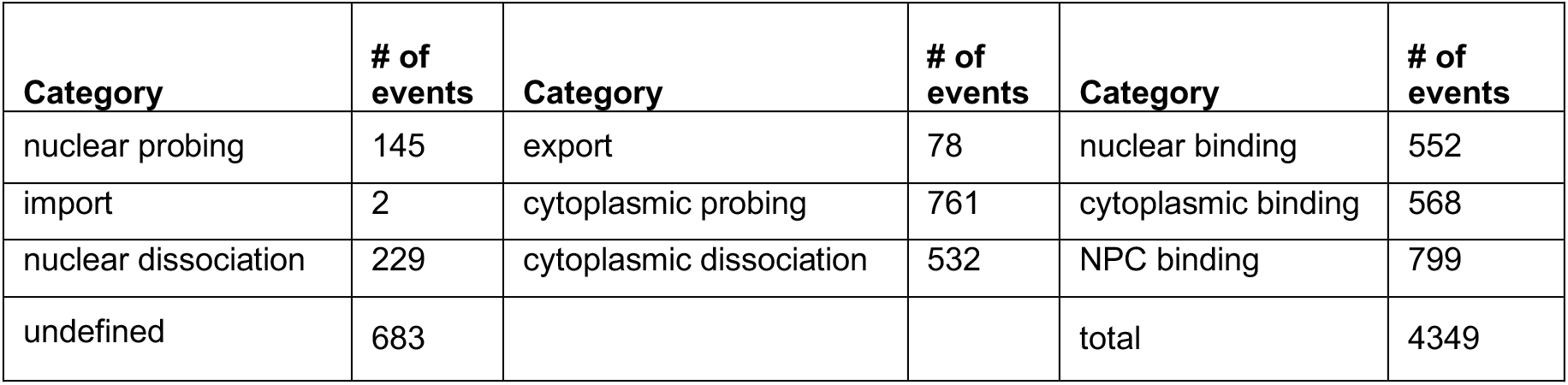
Number of detected interaction events with single NPCs sorted according to the type of interaction.

| Category | # of events | Category | # of events | Category | # of events |
| --- | --- | --- | --- | --- | --- |
| nuclear probing | 145 | export | 78 | nuclear binding | 552 |
| import | 2 | cytoplasmic probing | 761 | cytoplasmic binding | 568 |
| nuclear dissociation | 229 | cytoplasmic dissociation | 532 | NPC binding | 799 |
| undefined | 683 |  |  | total | 4349 |

### Abortive export events

In addition to the complete observed export events our analysis also returned attempted or abortive export events in the category “nuclear probing” (Fig. 3G). These were defined by a particle approach to the NE from the nucleoplasm, an interaction with a specific NPC, and then return back into the nucleoplasm. Interestingly, in this category we observed more events, namely 145 (Table 1). Obviously, also the category “nucleoplasmic dissociation” comprising 229 processes described failed or interrupted export events.

Again, we determined the interaction duration of the failed export events with the NPCs and the distribution of binding sites along the NPC symmetry axis (Fig. 5). Not surprisingly, the interaction duration of the subunits, whose export failed, was shorter than that of completed exports, namely only τ=13±3ms.

## Discussion

Here, we report on the nuclear export of single pre-ribosomal particles through NPCs by live microscopy. The analysis of the transport process kinetics and of the binding site distribution of the pre-60S particles along the transport axis of NPCs allowed us to define key steps of the export process and the position of the rate-limiting step of the translocation through the central NPC channel.

In order to visualize single pre-60S particles we used an indirect fluorescence labeling approach. We stably expressed eIF6-HaloTag in HeLa cells and labeled it in vivo by the fluorescence dye JF549. Using extremely low fluorophore concentrations it was possible to observe single, mostly mobile, diffraction limited spots in live cells by sensitive fluorescence microscopy. We provided five-fold evidence that these spots represented pre-60S particles. In bulk experiments, the fluorescent particles showed a nucleolar accumulation mirroring the localization of tagged ribosomal proteins, such as RPL29, in pre-60S reporter cell lines^18^. Addition of the transcriptional inhibitor actinomycin D resulted in a loss of the typical nucleolar fluorescence, presumably caused by a stop of the production of pre-ribosomal particles. A pull-down experiment demonstrated that the labelled eIF6 was contained in molecular aggregates that comprised also the ribosomal protein Rpl10, an established component of pre-60S particles(reviewed by ^42^). Incubation with LMB, an inhibitor of pre-60S nuclear export by blocking Crm1, resulted accordingly in an increase of nuclear fluorescence. Finally, the observed single particles diffused more slowly than the smaller pre-40S particles and drastically more slowly than unbound proteins. Altogether this suggested that eIF6-HaloTag-JF549 was contained in pre-60S particles and allowed us to visualize pre-60S particle dynamics in living cells.

We used super resolution confocal laser scanning microscopy of eGFP-NTF2-expressing HeLa cells to reveal the location of the NE. Distinct and characteristic fluorescence maxima along the NE indicated the positions of single NPCs. Quasi-simultaneously we imaged the nuclear transport of the pre-60S subunits in the same cellular region. In this way we were able to observe pre-60S particles interacting with single NPCs. These events were analyzed by an elaborate automatic data processing pipeline.

Altogether, we identified 78 complete export events for which we monitored an approach from the nuclear side to the NPC, an encounter with the NPC and a release into the cytoplasm. The data analysis extracted the transport trajectories during this process revealing interaction durations (dwell times) with the NPCs and the respective binding positions. The distribution of the dwell times at the NPCs could best be described by a monoexponential decay function as was shown by the analysis of the distribution using the Akaike information criterion yielding an expectation value for the translocation duration of 24±4 ms.

This value is in very good agreement with estimations for the maximum transport rate of cargo molecules through the human NPC formerly estimated by Ribbeck and Görlich (2001, 2002)^33,43^. From the electron tomography study of Delavoie et al (2019) it is evident that ribosomal subunits are exported one at a time and not in parallel. Thus, the translocation duration of 24±4 ms in principle would allow the transport of ~35 to 50 large subunits per second through a single pore. Assuming an average molecular mass of ~2.5 MDa for the human pre-60S particle this accounts for a mass flux of 87.5 to 125 MDa·s^-1^ per NPC. Our live cell data show that the maximum transport rate is almost reached in vivo and is higher than the suggested in-vivo minimum rate of 10-40 MDa·s^-1^ for a growing cell^22,43^.

Notably, the measured translocation duration of 24±4 ms is in the same order of magnitude as the ~90±50 ms for the dwell time of pre-60S particles in the yeast NPC, which was estimated by Delavoie et al. (2019) ^23^. These authors observed pre-60S particles within NPCs in EM tomograms and used a probabilistic queueing model to extract a transport dwell time from the NPC occupancy. Among other parameters their estimate was based on the number of NPCs available for transport. Smaller NPC numbers would lead to shorter translocation times. In our experiments we observed – apart from complete export events – also a large fraction of attempted or “failed” export events. These unsuccessful events might indicate that numerous NPCs are temporarily ‘occupied’, e.g. with other bulky substrates and indeed not accessible for ribosomal export. This would effectively reduce the number of NPCs and lead to a lower translocation time estimate in the queueing model. Therefore, we consider our result as a support of their model.

Ribbeck and Görlich (2002) found that the transport rates of bigger transport cargoes scale with the number of loaded transport receptors^43^. This was later also shown for exceptionally big, artificial transport substrates, namely quantum dots (QD) functionalized with importin β binding domains featuring a hydrodynamic radius of ~18 nm^44^. Reducing the number of bound importin β molecules dramatically increased the QD dwell time in the central NPC channel^44^. Intriguingly, in yeast at least eight different transport receptors are loaded on a single pre-60S particle, i.e. Mex67/Mtr2, Rrp12p, Arx1, Ecm1, Bud20, Npl3, Gle2 and Xpo1, but for the human pre-60S particle so far only two main export receptors are known: Crm1 and Exp5. Only one Crm1 can be loaded to the NES of NMD3. Exp5 can recognize structurally diverse RNAs^22^ and recognizes its rRNA cargo in a sequence-independent manner via double-stranded stem-loops^18^. It is tempting to speculate that in this way eventually more than one Exp5 is loaded on a single human pre-60S particle to ensure the necessary ‘receptor density’ for such a bulky cargo.

After the translocation the cargo/receptor complex is dissociated on the cytoplasmic side of the NPC. Mex67/Mtr2 is removed by the ATP-dependent helicase activity of Dbp5 from mRNA, but surprisingly this does not occur for pre-ribosomal export^45^. Also, for the non-canonical export factors in yeast, e.g. Arx1, it is unknown how and when they are detached from the yeast pre-60S particle. In contrast, for the RanGTPase-dependent receptors Exp5 and Crm1 engaged in human pre-60S particle export it is known that they are released after RanBP1/RanGAP1-triggered GTP hydrolysis. For Crm1-mediated export, nucleoporin Nup214 has been shown to play a critical role in this final step ^46,47^. Moreover, Nup214 (Nup159 in yeast) is also critical for the pre-60S particle export in mammalian^18^ and yeast cells^48^. RanGAP cannot directly act on Ran in the export complex, but needs RanBP1 or the Ran-binding domains of RanBP2/Nup358^22^. Nup214, Nup88 and Nup358 are the only nucleoporins located asymmetrically on the mammalian NPC and in human cells the Nup214/Nup88 sub-complex mediates the attachment of Nup358 to the NPC^49^. Our finding of a single rate limiting step and single maximum of binding sides in the central NPC suggests a model, where both Exp5 and Crm1 are needed for NPC entry and passage of the pre-60S particle, but only the Nup214/Crm1 interaction is needed for release from the NPC. Here, the strong and direct interaction between Nup214/Crm1 would ‘fish-out’ the pre-60S export complex from the central NPC channel via FG-repeats^50^ and hand it over to NUP358, which results in GTP-hydrolysis and subsequent cytosolic pre-60S release (supplemental Fig. 11).

Our analysis of the binding site distribution of subunits along the translocation axis and the topology of NPCs (Fig. 5) confirmed the results of Delavoie et al. (2019) and allowed a comparison with mRNP export. Our measured distribution of trajectory positions, although less well resolved than the EM data, showed a clear, single maximum in the central region of the NPC shifted towards the cytoplasm. Delavoie et al. (2019) observed a relatively low fraction of pre-ribosomes in the region of the nuclear ring, a large fraction at the inner ring of the NPC, and a medium-scale fraction at the cytoplasmic ring of the NPC, i.e. their binding site distribution showed a single maximum and a shoulder near the cytoplasmic filaments^23^. In agreement with their data, we also did not see an accumulation of binding sites in the region of the nuclear basket. Delavoie et al. (2019) suspected therefore that the pre-60S particle-nuclear basket interactions are of very transient nature^23^. This was supported by our observation that the dwell time of “nuclear probing” (13±3ms) was short compared to the actual translocation step (24±4ms). The reason might be the missing quality control step for pre-ribosomes at the nuclear basket^23,51,52^, like it exists for mRNPs^53,54^.

Interestingly, for mRNP export trajectories a complementary distribution with two maxima on the nucleoplasmic and cytoplasmic faces of the NPC were observed^41,55^. It was suspected that the maximum at the nuclear face is due to quality control processes and/or RNP reorganization for the translocation, and the maximum at the cytoplasmic face due to the removal of the mRNP export factors by the RNA helicase Dbp5, which infers the directionality of mRNP export (reviewed by ^56^). As already stated, such complex enzymatic processes do not occur for pre-ribosomes.

Our data analysis allowed to discriminate between successful and failed transport events. In the latter case, the respective pre-60S particles also interacted with NPCs according to our definition. Interestingly, we observed this significantly more often (n=145) than successful export processes (n=78). Thus, only about 35% of attempted export events were successful. Similar fractions have been reported for mRNPs, so the value of about 1/3 of successful export events may present a general rule for large substrates. The binding site distribution of the failed exports showed a distinct maximum in the region of the nuclear basket, just outside the nuclear ring of the NPC. This supported the view that the respective pre-60S particles did not enter the hydrophobic NPC interior, but were blocked from it. We speculate that this was either due to an occupancy of the NPC by another large transport cargo or indicated that the particles still missed an export factor, possibly Crm1.

In conclusion, we showed that nuclear export of pre-60S particles takes 20 to 30 ms and that about 1/3 of attempted export trials are successful. The multi-step light microscopic approach introduced here is laborious, but allowed for the first time the observation of single pre-60S particles during their interaction with NPCs in vivo. For the future, it is important to increase the experimental throughput. In that case it could be combined with the mutational or RNA interference analysis of the nuclear transport machinery to refine our understanding of the transport of large RNA cargoes such as pre-ribosomes through the NPC. A comparable approach to study the nuclear export of pre-40S particles appears straightforward and would extend the range of questions to be asked.

## Acknowledgements

We gratefully acknowledge a grant of the Swiss National Science Foundation (SNSF) to Ulrike Kutay (31003A_166565 and NCCR ‘RNA and disease’).

