## Supplemental Figures for "Nuclear export of the pre-60S ribosomal subunit through single nuclear pores observed in real time"

### Supplemental Material

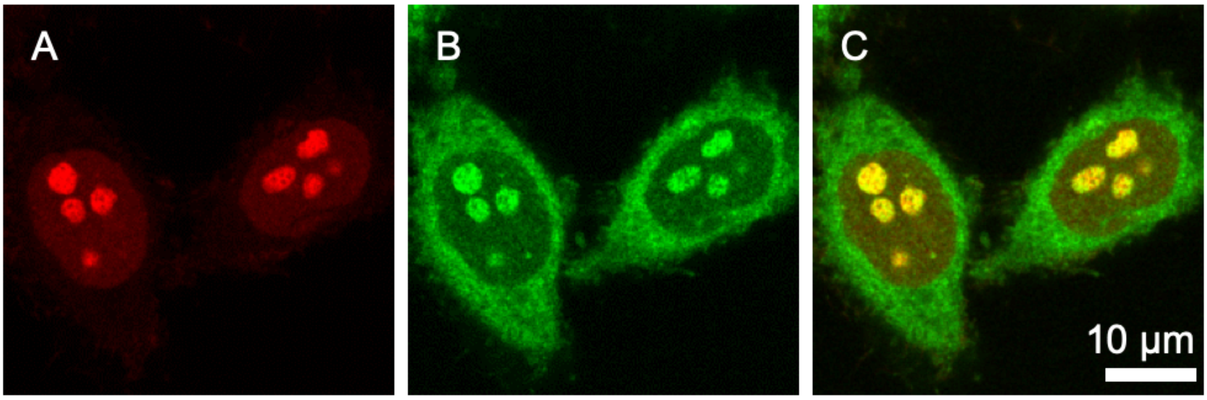


**Supplemental Figure 1**

eIF6-HaloTag-JF549 shows the same nucleolar distribution as Pno1-Snap-SiR647[1,2]. (A) HeLa cells expressing Snap-Pno1 stained with 647-SiR. (B) Transiently expressed eIF6-HaloTag stained with JF549. (C) Overlay of both channels.

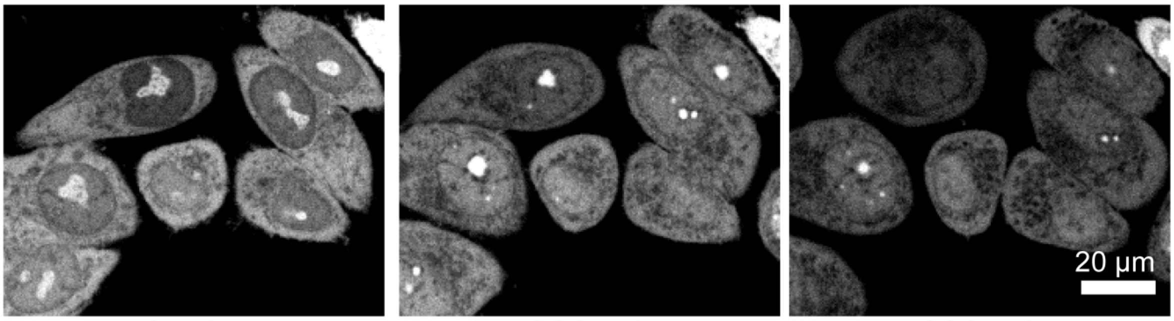


**Supplemental Figure 2**

Actinomycin D inhibits synthesis of new transcripts and leads to a loss of nucleolar staining implying that no further subunits were formed. Confocal sections of HeLa S3 cells stably expressing eIF6-HaloTag labelled by JF549 upon addition of 10 mM Actinomycin D. Images were taken subsequent to Actinomycin D addition: (A) 0 min, (B) 300min, (C) 600 min.

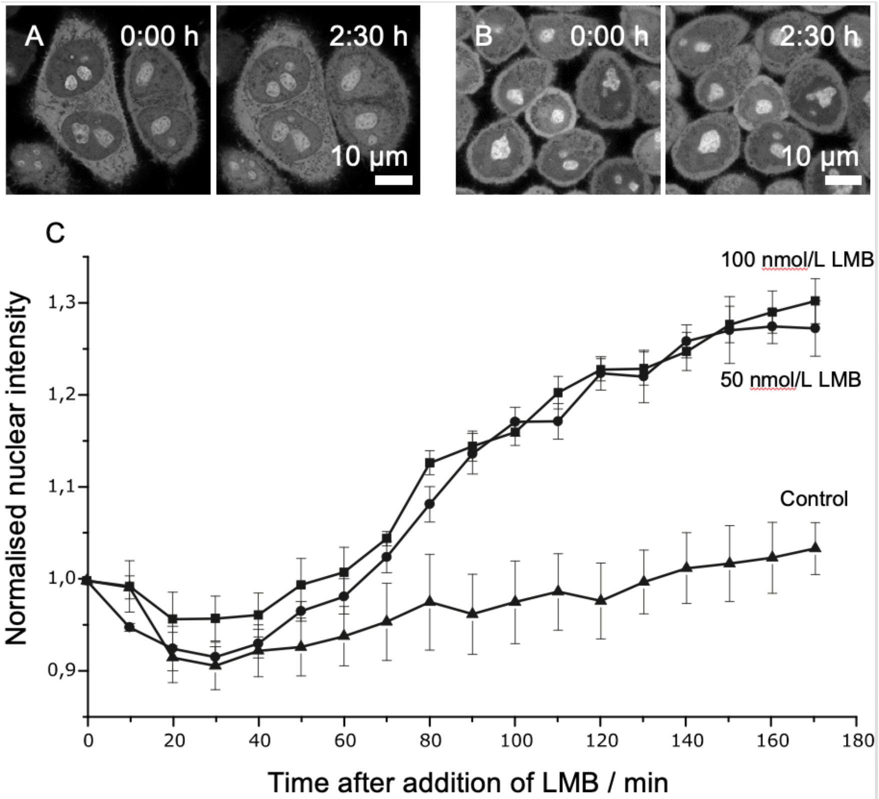


**Supplemental Figure 3**

LMB prevents Crm1 binding and thus nuclear export of the pre-60S subunit leading to an increase of intranuclear fluorescence. (A) eIF6-HaloTag-JF549 at 0:00 h and after 2:30 h incubation with 100 nmol/L LMB. (B) eIF6-HaloTag-JF549 at 0:00 h and after 2:30 h incubation with 10 µL ethanol as control (C) Normalised fluorescence intensity in the nucleus as a function of time at a concentration of 50 nmol/L LMB (N=7, circles), a concentration of 100 nmol/L (N=9, squares) and a control measurement with ethanol (N=4, triangles). Error bars represent the standard error of the mean.

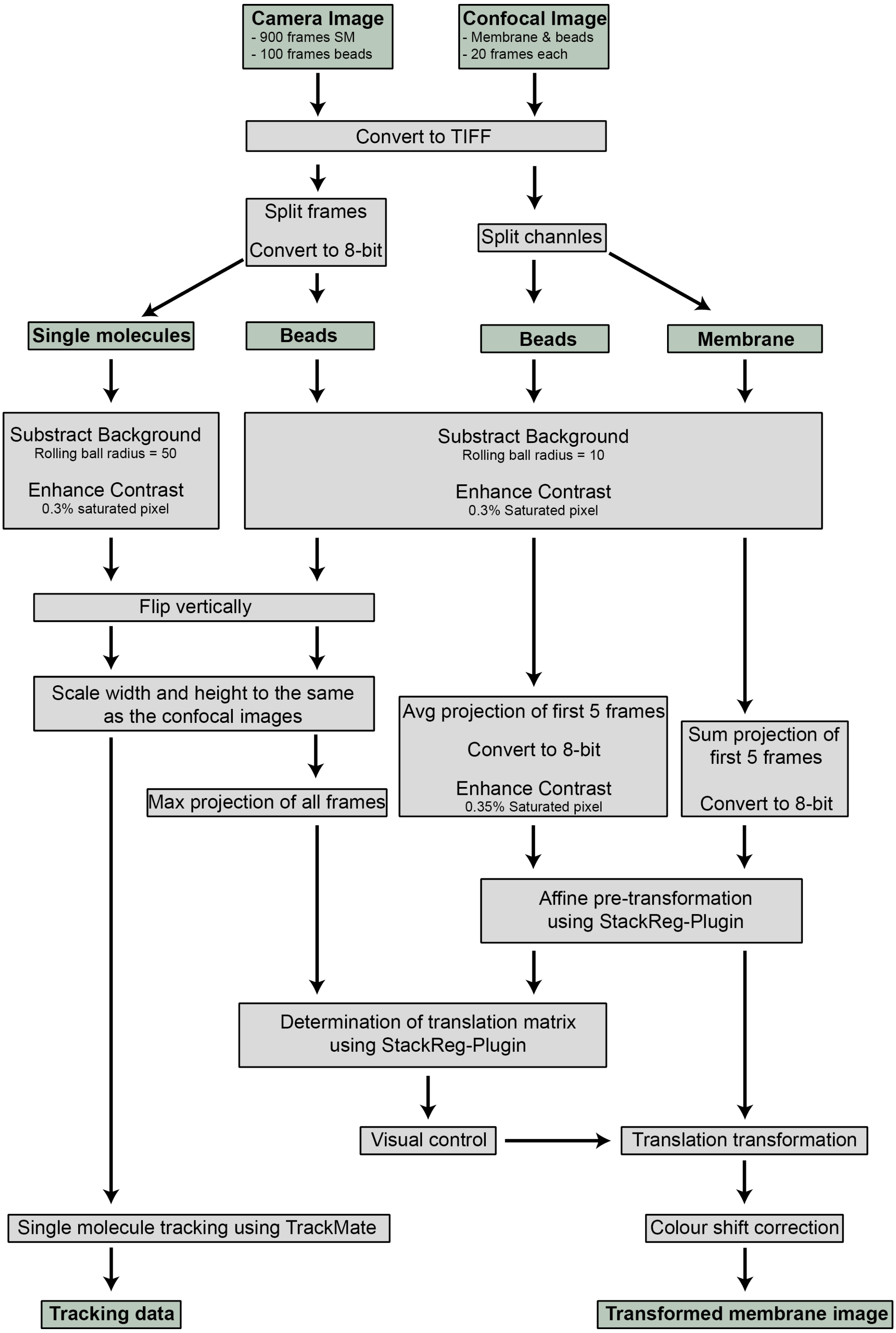


**Supplemental Figure 4**

Registration of images showing pre-60S particles and nuclear envelopes acquired with an EMCCD and a LSM 880 Airyscan, respectively. First, a movie of 1000 frames was acquired by the EMCCD camera. 900 frames were imaged with 561 nm excitation revealing the pre-60S subunits and 100 frames with 405 nm showing the reference beads. Next, 20 images were acquired using the LSM 880 Airyscan with alternating laser excitation (405 nm and 488 nm). Image processing of the EMCCD camera images: Image sequences were converted '8-bit TIFF', split into 900 frames of pre-60S subunits and 100 frames reference beads. Both image stacks were background-subtracted, contrast-enhanced and vertically flipped. A maximum intensity projection was calculated from the reference bead stack. Single particles were tracked using TrackMate plugin for ImageJ. Image processing of the confocal images: Airyscan images were processed and converted into TIFF format. The two channels were separated, background-subtracted and contrast-enhanced. The first five frames of the reference beads were averaged and the contrast was again enhanced. The first five frames of the membrane image were summed. Both projections were converted to 8-bit. To align the EMCCD camera and Airyscan images, an affine transformation was applied using the StackReg plug-in for ImageJ to correct for rotation, shear and – roughly - the translation. To correct for stage drift during the measurement a further transformation matrix was determined from the bead images and applied to the Airyscan image of the NE. Finally, images were colour shift corrected.

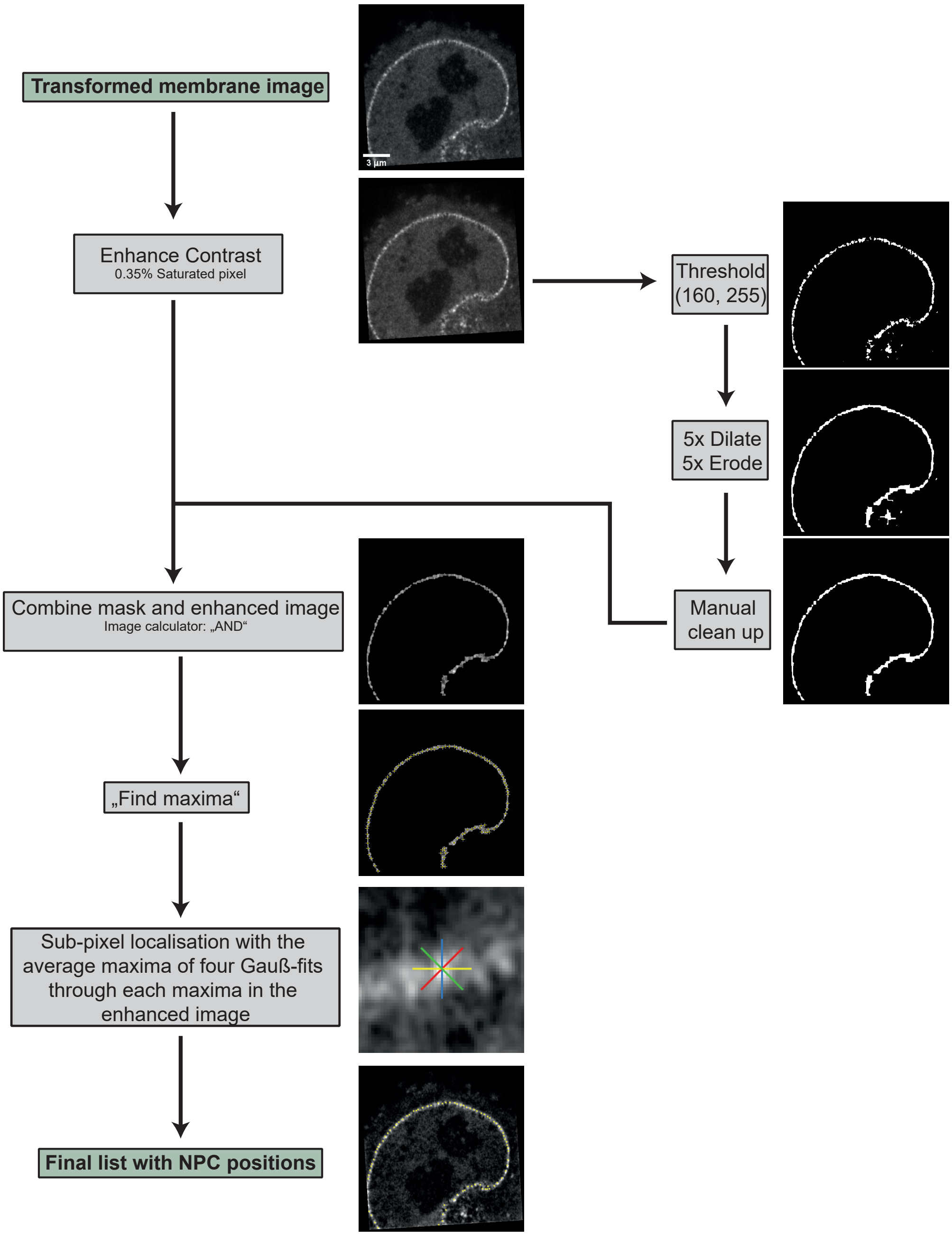


**Supplemental Figure 5**

Determination of single NPC positions. For details, see Methods.

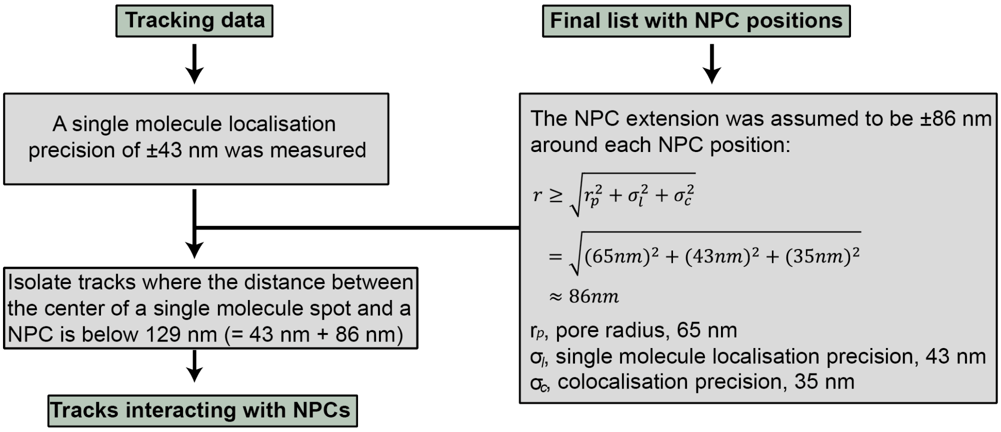


**Supplemental Figure 6**

Identification of single pre-60S particle tracks, which interact with single NPCs. For details, see Methods.

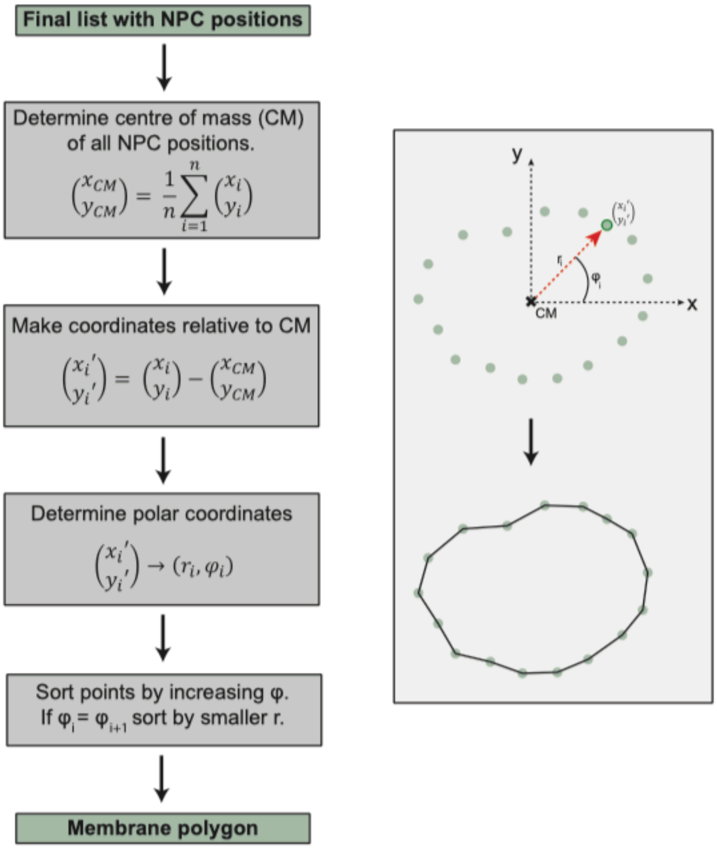


**Supplemental Figure 7:**

Derivation of the polygon, which approximated the shape of the cell nucleus. For details, see Methods.

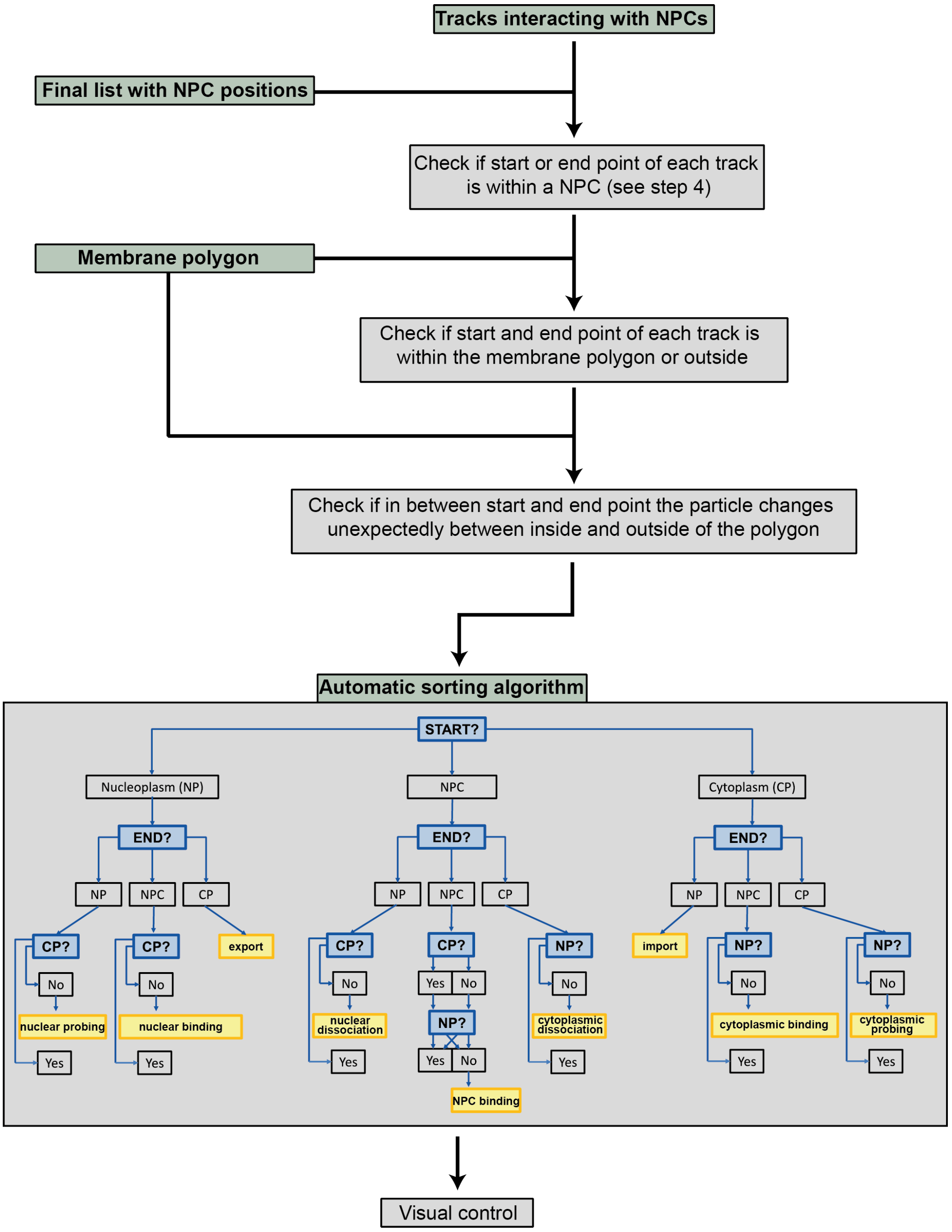


**Supplemental Figure 8**

Automated sorting of the observed trajectories interacting with single NPCs into the various contact modes as defined by Fig. 3G. For details, see Methods.

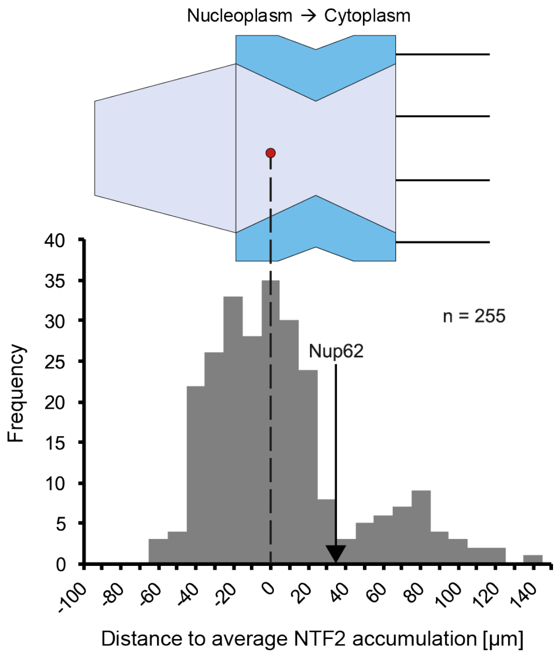


**Supplemental Figure 9**

Maximum NTF2 fluorescence intensity as indicator for the NPC center location, which was calibrated using the anti-nucleoporin p62 (Nup62) antibody mAB414 as reference. For 255 NPCs in 33 cells the positions of the HaloTag-NTF2-JF549 and Nup62 (stained using mAb414 and a polyclonal anti-mouse secondary antibody labelled by AlexaFluor 488) maxima were determined by fitting four Gaussian functions in 45° angles (see supplemental Fig. 5). The maxima of the fits were averaged and the distance between the maxima of the Nup62 and NTF2 was calculated. The final distances were corrected by 10 nm to account for the Nup62 signal position with regard to the central plane of the NPC as determined by [3,4] and shifted the whole histogram so that the average NTF2 maximum was at position 0. The average distance of NTF2 from the central plane of the pore is about -24 nm. Interestingly, we observed two maxima of the binding of NTF2, as has been observed previously[5–7].

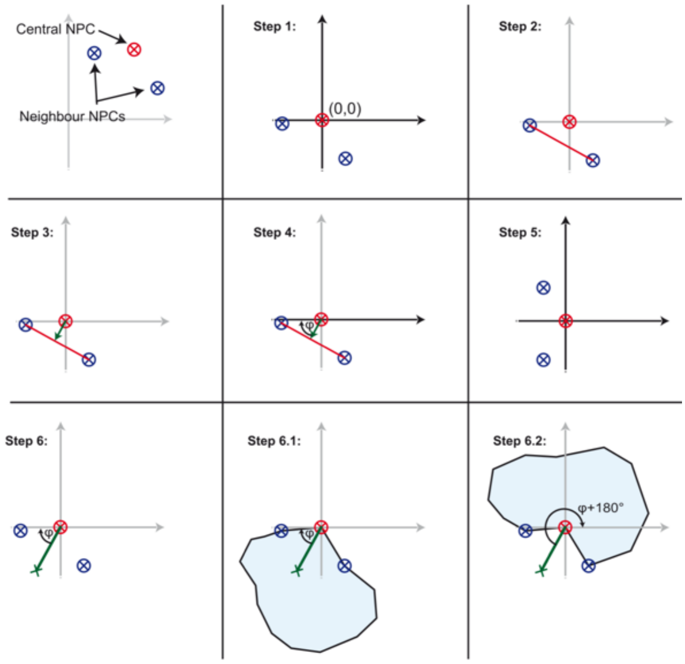


**Supplemental Figure 10**

Alignment of the export tracks onto the origin and the export direction to the abscissa. The coordinate system was shifted such that the transporting, "central" NPC (marked in red) represented the new origin. We assumed that export occurred along the axis, which was perpendicular to the connecting line between the two NPCs (blue marks), which were neighboured to the transporting, "central" NPC (marked in red). This line was parallel to the ordinate in Step 5. Usually, the NPCs formed a convex line (Step 6.1), but in rare cases the NE line was concave, which was considered by rotating the track coordinates by φ+180° (step 6.2).

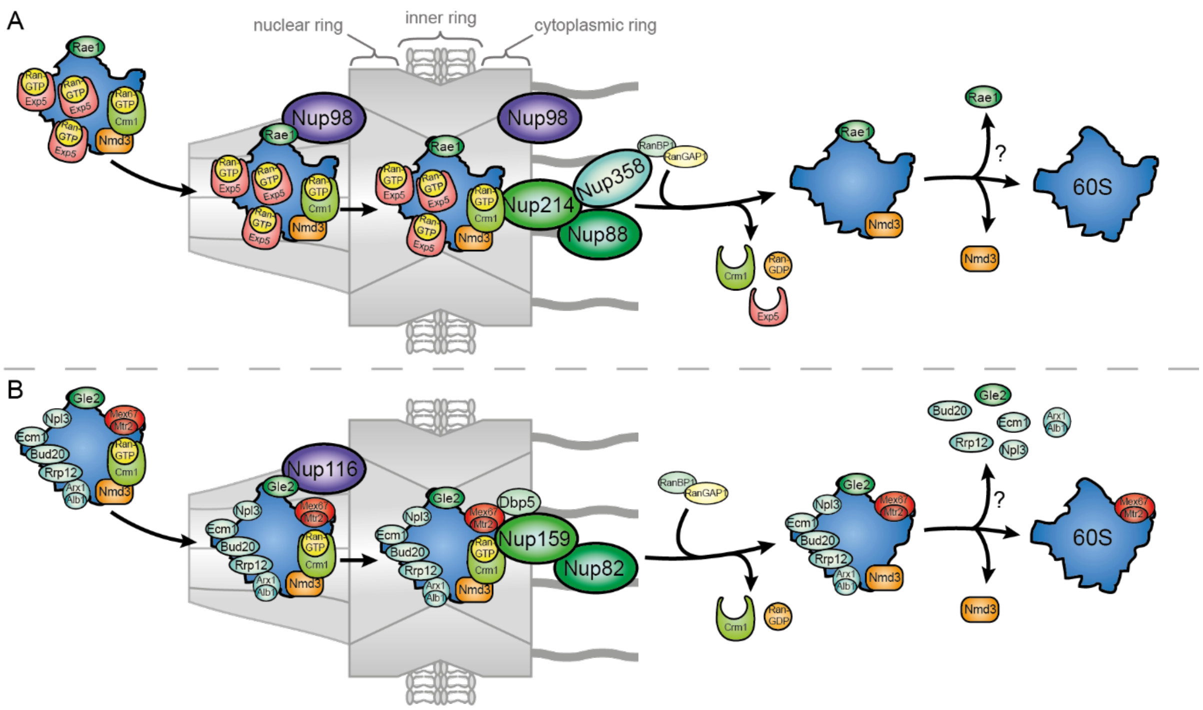


**Supplemental Figure 11**

Proposed pre-60S particle export mechanism in (A) human cells. Human pre-60S export begins with the non-FG interaction of RAE1 with Nup98. The entry into the FG-repeat domain of the NPC channel is mediated by Exp5 and Crm1. Release from this requires the pre-60S particle/CRM1/Nup214 interaction at the cytoplasmic face of the NPC, which “fishes” the particle out of the central NPC domain. Subsequently the pre-60S particle escapes into the cytosol after CRM1 release induced by RanBP1/RanGAP1, which is recruited to the NPC by Nup358. (B) In yeast the receptor-loaded pre-60S particle first transiently interacts with Nup116 at the nuclear basket via Gle2 before being transported into the inner ring in a Mex67/Mtr2-dependent manner. Release from this energetically favourable NPC compartment would then similarly require 'fishing out' the Crm1/pre-60S particle cargo through strong interactions with the asymmetric NUP159, which is part of the Nup82 subcomplex, and subsequent removal of Crm1 by RanBP1/RanGAP1 for cytoplasmic release of the pre-ribosomes.

**Supplemental Movie S1**

HeLa cell stably expressing GFP-NTF2 and eIF6-HaloTag labelled with JF549-HaloTag-Ligand. Single pre-60S particles (red) diffuse within a nucleus. The nuclear envelope is shown in green. Scale bar, 3 µm.

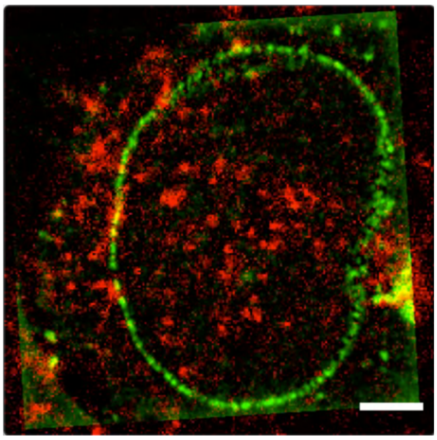

**Supplemental Movie S2**

Single pre-60S particle (red) passing the nuclear envelope (green). The image data was smoothed and contrast enhanced to improve the visualization. Scale bar, 3 µm.

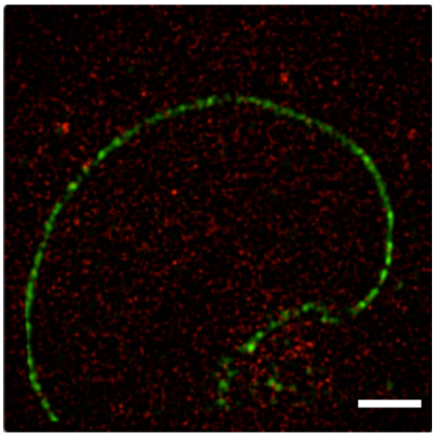
